# Histology-informed spatial domain identification through multi-view graph convolutional networks

**DOI:** 10.1101/2025.10.23.683862

**Authors:** Huihui Zhang, Jiaxing Chang, Yue Sun, Pinli Hu, Jing Liu, Jiuju Luo, Hang Yang, Yonglin Ren, Xingtan Zhang, Zehua Chen, Kok Wai Wong, Haojing Shao

## Abstract

Identifying spatial domains is crucial in spatial transcriptomics, yet effectively integrating gene expression, spatial location, and histology remains challenging. We present STESH, a Spatial Transcriptomics clustering method that combines Expression, Spatial information and Histology. STESH extracts histological features using a convolutional neural network and generates expression, histology, spatial, and collaborative convolution modules for a multi-view graph convolutional network with a decoder and attention mechanism. We evaluated STESH on multiple tissue types and technology platforms. STESH consistently outperformed ten state-of-the-art methods, achieving superior clustering accuracy with the highest scores in adjusted Rand index, normalized mutual information, and Fowlkes-Mallows index.

## Introduction

Identifying the spatial domain is the first important step in spatial transcriptomics (ST). Currently, methods for identifying spatial domains can be roughly divided into three categories by the number of dimensions: one dimension (expression) clustering, two dimensions (expression and spatial) clustering, and three dimensions (expression, spatial and histological) clustering. Expression only clustering methods, such as k-means and Louvain^1^, focus solely on gene expression data, and ignore spatial and histological contexts, which leads to discontinuous domain clustering. To address this, expression and spatial clustering methods such as Giotto’s Hidden Markov Random Field model, SEDR’s ^2^ deep autoencoder network, and STAGATE’s ^3^ attention mechanism incorporate spatial information to improve domain identification. Other methods, like CoSTA^4^, BASS^5^, CCST^6^, BayesSpace^7^, GraphST^8^, SCGDL^9^, and AE-GCN ^10^, use various strategies, including convolutional neural networks, Bayesian approaches, and graph neural networks to enhance clustering accuracy by integrating spatial and gene expression data. These methods jointly utilize gene expression and spatial localization but do not exploit the histological information that high-resolution histological images can provide.

Expression, spatial and histological methods extract features from histological images to assess histological similarity. Combined with the spatial neighborhood structure, this similarity is then used to refine gene expression. SpaGCN^11^ represents a pioneering graph convolutional network (GCN) algorithm that integrates histology, spatial coordinates, and gene expression data. This approach constructs an undirected weighted graph by combining histology images, spatial coordinates, and gene expression profiles, which is subsequently incorporated into the GCN framework. stLearn^12^ combines histological features from histological images with gene expression from neighboring spots to achieve domain clustering. TIST^13^ converts hematoxylin and eosin (H&E) images to grayscale and extracts image features based on a Hidden Markov Random Field (HMRF) model. This image processing strategy results in the loss of color information, hindering the consideration of highly correlated color pixels. DeepST^14^ uses a pre-trained deep neural network model to extract feature vectors from histological image tiles, then integrates the extracted features with gene expression and spatial location data to characterize the correlation of spatially adjacent spots and create a spatially enhanced gene expression matrix. These methods establish a connection between image features and gene expression data but do not fully involve image information in subsequent training, which may lead to inaccuracies.

Integrating histological images, gene expression data, and spatial location information offers a promising approach for spatial clustering of spatial transcriptomics (ST) data. However, existing methods struggle to incorporate spatial information and align with high-resolution histological images. In this study, we developed a novel framework to effectively utilize histological images for spatial domain identification. Specifically, we employed a state-of-the-art convolutional neural network (CNN) to extract and learn detailed texture features from histological images, constructing corresponding graphs. To ensure the independence of histological patterns from spatial and expression information, we constructed three separate GCN for spatial, histological, and expression data, respectively. Furthermore, we added a fourth integrative GCN that collaboratively learns from all three data types to capture the intrinsic relationships among these modalities.

## Result

### Overview of STESH

The main workflow of STESH is a multi-view graph convolutional network in deep learning. The first step is constructing three adjacent graphs for histological information from images, spatial information, and gene expression (Figure 1A). The histological graph is calculated by pretraining convolutional neural networks method such as ResNet50 (Figure 1B)^15^. The next step is reducing the dimension by autoencoder and decoder (Figure 1C). The last step is explaining the low-dimensional data by spatial clustering and trajectory inference (Figure 1D).

**Figure 1.**
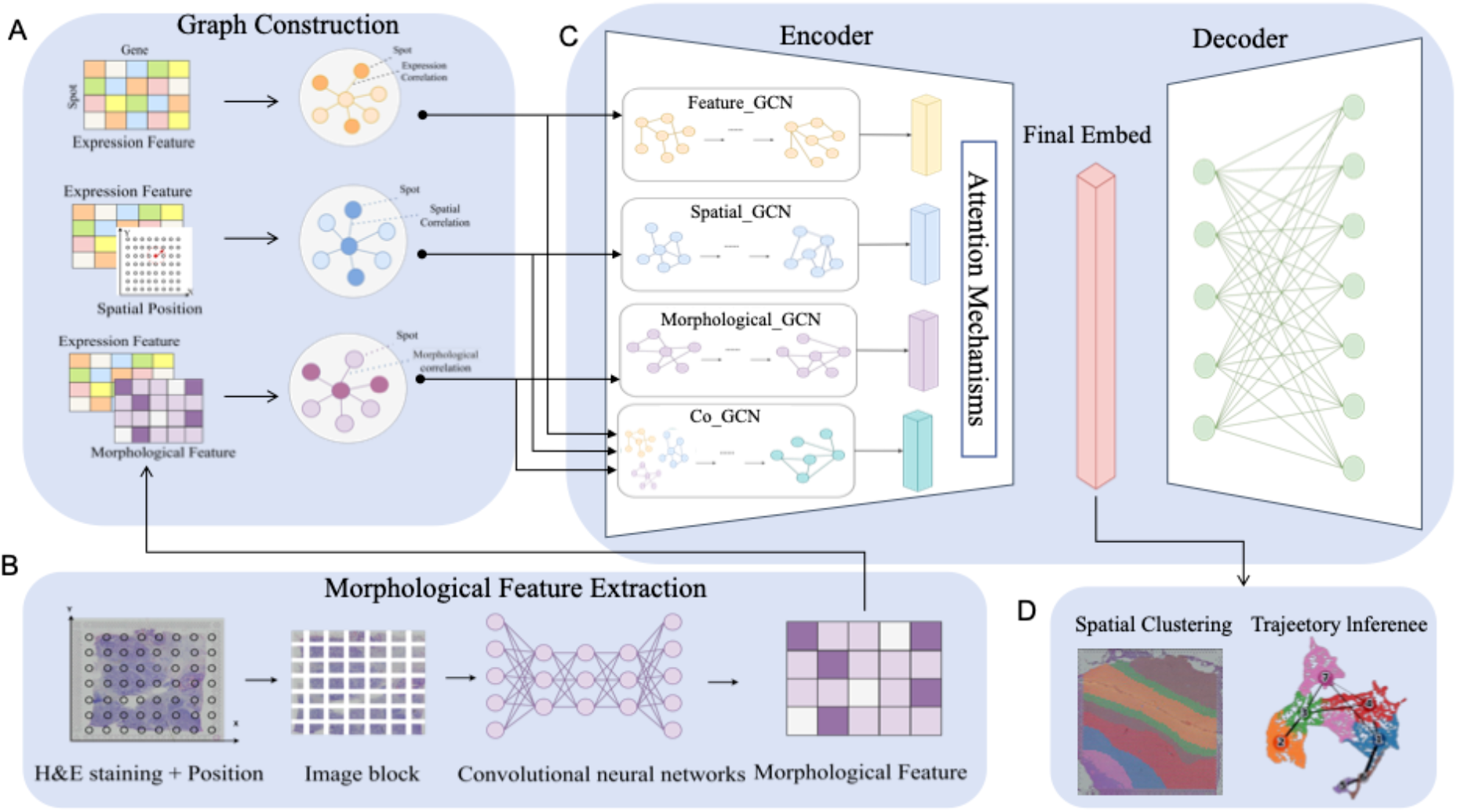
Workflow of STESH algorithm. **A**, Graph Construction. Based on the measured gene expression data, spatial location information and tissue section images in the spatial transcriptome, the spatial adjacency matrix, the expression adjacency matrix and the morphological adjacency matrix were constructed through different similarity measures. Combined with the gene expression feature, the expression graph, the spatial graph and the morphology graph were constructed. **B**, Morphological Feature Extraction. STESH utilizes H&E staining to extract histological features. Initially, the center of the image patch is determined based on the actual coordinates of the spot within the section. Subsequently, the tissue section image is cropped into square image patches of an appropriate size. Then, a pre-trained convolutional neural network (CNN) model is employed as the feature extractor, taking the segmented image patches as input to extract morphological information from the tissue section images. **C**,Multi-view GCN encoder. The GCN encoder uses feature GCN, spatial GCN, Morphological GCN,and Co-CCN, to learn the latent embeddings of spots on the feature graph, spatial graph and morphology graph, and adaptively fuses these embeddings with the learned weights using an attention mechanism. **D**, Application. Applying learned low-dimensional embeddings to spatial clustering and trajectory inference.

For benchmarking STESH, we utilized three public datasets that were widely used for comparison [SEDR, DeepST, STAGATE, stLearn]. The human dorsolateral prefrontal cortex (DLPFC) dataset contains 12 sections (3460-4789 spots), which were manually annotated in seven cortical layers. The human ovarian cancer dataset (3493 spots) is also manually annotated in 20 layers. These two datasets are generated by 10X platform. To demonstrate that STESH can be applied to high-resolution data, we used a mouse olfactory bulb dataset (19109 spots) generated by Stereo-seq^16^.

To quantitatively evaluate the contribution of histological image integration in STESH, we developed an image-free version of our framework. Comparative analysis revealed that the image-integrated GCN architecture significantly enhances clustering accuracy, demonstrating the substantial value of histological information in spatial transcriptomics analysis(Supplementary Figure 1). STESH uses a fixed random seed (seed=100) throughout its implementation and our experiments (Supplementary Figure 2).

In order to comprehensively and systematically evaluate the performance of the spatial domain recognition method proposed in this article, ten representative spatial domain recognition methods were selected as comparison benchmarks. Leiden is a non-spatial method widely used and implemented in Seurat ^17^ and Scanpy. GraphST, Spatial-MGCN, BayesSpace, SpaceFlow, SEDR, and STAGATE are spatial methods that do not require histological images, while stLearn, SpaGCN and DeepST are spatial methods that use histological images. The spatial domain recognition performance of each method is measured using three common clustering metrics: the Adjusted Rand Index (ARI), Normalized Mutual Information (NMI), and Fowlkes-Mallows Index (FMI).

### Benchmarking STESH on DLPFC dataset

Our analysis revealed that all spatial methods outperformed the non-spatial method Leiden, demonstrating the significant contribution of spatial information to clustering accuracy. As illustrated in Figures 2A-C, STESH achieved superior performance with average ARI (0.61), NMI (0.69), and FMI (0.70), surpassing ten state-of-the-art methods. For detailed evaluation, we selected slice #151672, which exhibits a well-defined five-layer cortical structure (Figures 2D). STESH accurately delineated all five cortical layers, achieving an exceptional ARI of 0.81 - the highest reported value for this dataset among recently published methods, including DeepST, SEDR, GraphST, SpaGCN, and Spatial-MGCN. While most methods maintained clear cluster boundaries, Leiden and stLearn exhibited less distinct separations. Notably, both Spatial-MGCN and STESH correctly identified layer 3 as a single cluster, whereas other methods erroneously split it into two clusters. The UMAP and PAGA trajectory plots further confirmed the robustness of STESH, GraphST, and Spatial-MGCN, all of which captured the sequential development of cortical layers with well-organized topological structures(Figures 2E).

**Figure 2.**
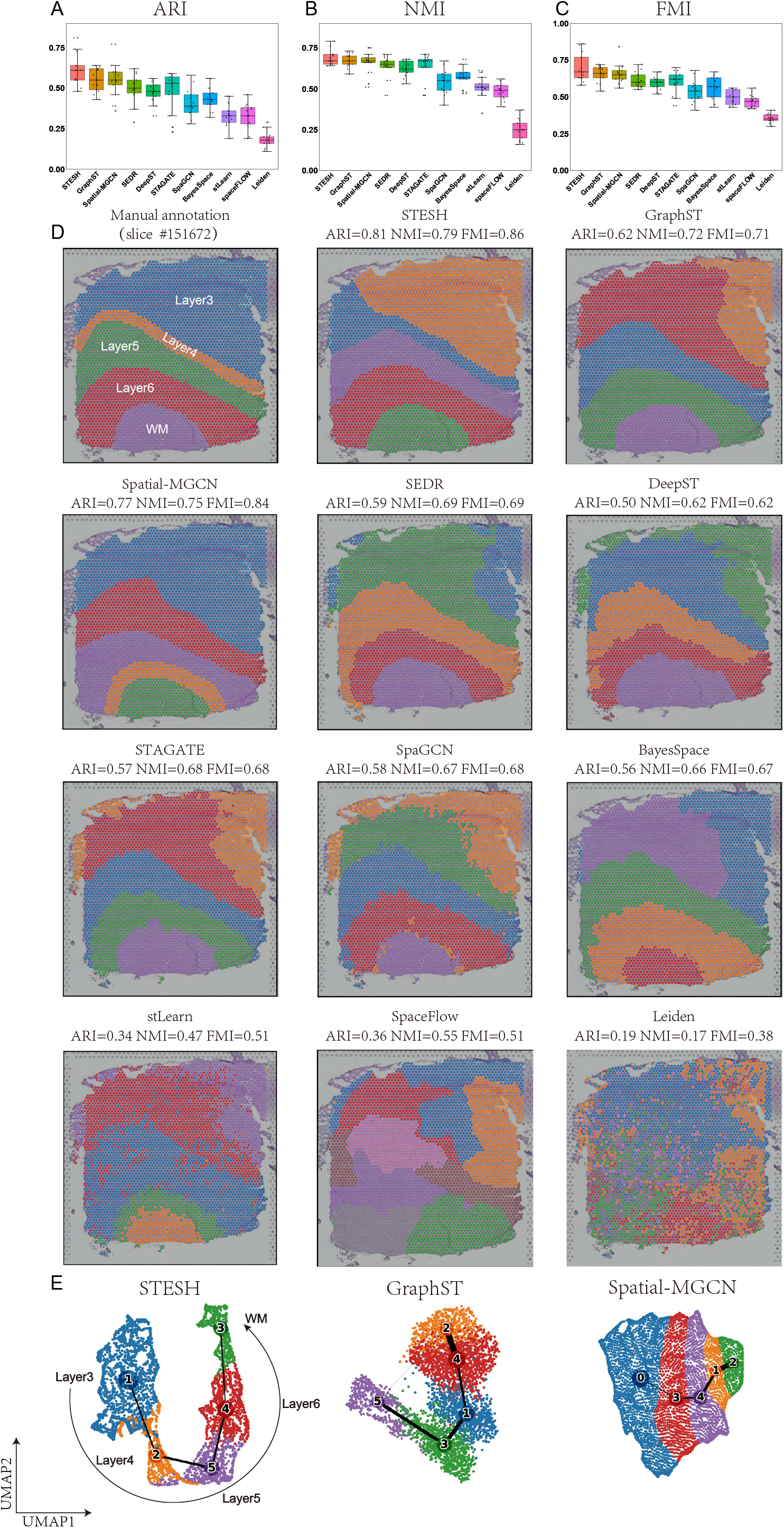
STESH improves spatial domain recognition in brain tissue. (**A**-**C**), Boxplot of the performance of STESH and other algorithms for all 12 DLPFCs. The y-axes are adjusted rand index (ARI), normalized mutual information (NMI), and Fowlkes-Mallows index (FMI) for **A, B**, and **C**, respectively. **D**, DLPFC layers of slide 151672 were annotated by Maynard^18^ et al., and identification of spatial domains by STESH, GraphST, Spatial-MGCN, SEDR, DeepST, STAGATE, SpaGCN, BayesSpace, stLearn, spaceFLOW, and Leiden. **D**, Trajectory inference and visualization of dimension reduction by Uniform Manifold Approximation and Projection (UMAP) for STESH, GraphST, and Spatial-MGCN, respectively.

**Figure 3.**
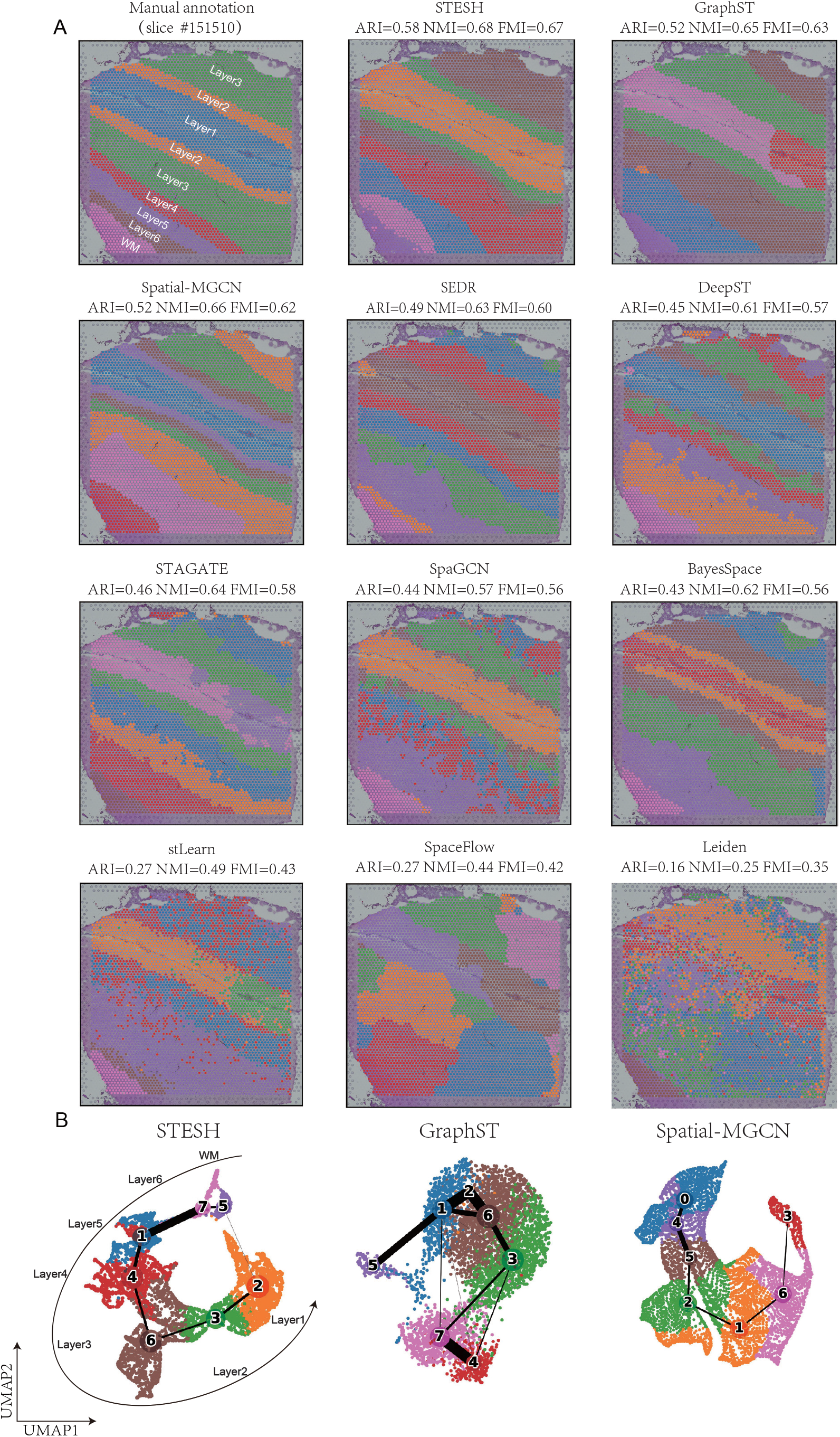
STESH improves spatial domain recognition in brain tissue slide 151510. **A**, The DLPFC slide was annotated by Maynard et al., and identification of spatial domains by STESH, GraphST, Spatial-MGCN, SEDR, DeepST, STAGATE, SpaGCN, BayesSpace, stLearn, spaceFLOW, and Leiden. **B**, Trajectory inference and UMAP visualization for STESH, GraphST, and Spatial-MGCN, respectively.

Slice #151510, characterized by a complex cortical structure comprising seven layers across nine regions, was selected for analysis. In this structure, layers 2 and 3 are divided into two regions flanking layer 1 (Figures 2A). STESH, GraphST, Spatial-MGCN, SEDR, BayesSpace, and SpaceFlow demonstrated well-defined cluster boundaries, whereas SpaGCN, stLearn, STAGATE, and DeepST exhibited discontinuous boundary delineation. Notably, GraphST, SEDR, BayesSpace, and SpaceFlow erroneously split layer 1 into two distinct clusters. Among all methods, only STESH and Spatial-MGCN successfully reconstructed the cortical structure with accuracy comparable to manual annotation. The UMAP and PAGA trajectory plots further revealed that both STESH and Spatial-MGCN exhibited a linear topological structure, precisely capturing the sequential development of cortical layers from Layer 1 to Layer 6 and the white matter (WM). In contrast, GraphST displayed an incorrect topological configuration, showing anomalous connections between Clusters (1, 2, 6) and Clusters (3, 7, 4), indicating potential misclassification of cortical layer identities. These results collectively demonstrate STESH’s superior capability in spatial domain identification and its ability to accurately reconstruct complex biological structures.

### Benchmarking STESH on a breast cancer dataset

Tumor heterogeneity detection represents another critical application of spatial transcriptomics clustering. We evaluated STESH and ten competing methods on a 10× Genomics Visium dataset of breast cancer, which was manually annotated into 20 distinct regions: four ductal carcinoma in situ/lobular carcinoma in situ (DCIS|LCIS) regions, eight invasive ductal carcinoma (IDC) regions, six tumor edge regions, and two healthy tissue regions. To ensure a fair comparison, all methods were constrained to identify 20 clusters, matching the manual annotation. STESH outperformed all other methods, achieving the highest clustering accuracy with an ARI of 0.61, NMI of 0.69, and FMI of 0.64 (Figure 4A). Notably, STESH, along with Leiden, stLearn, and STAGATE, correctly identified the IDC_4 tumor region as a single cluster without erroneous splitting, consistent with the manual annotation.

**Figure 4.**
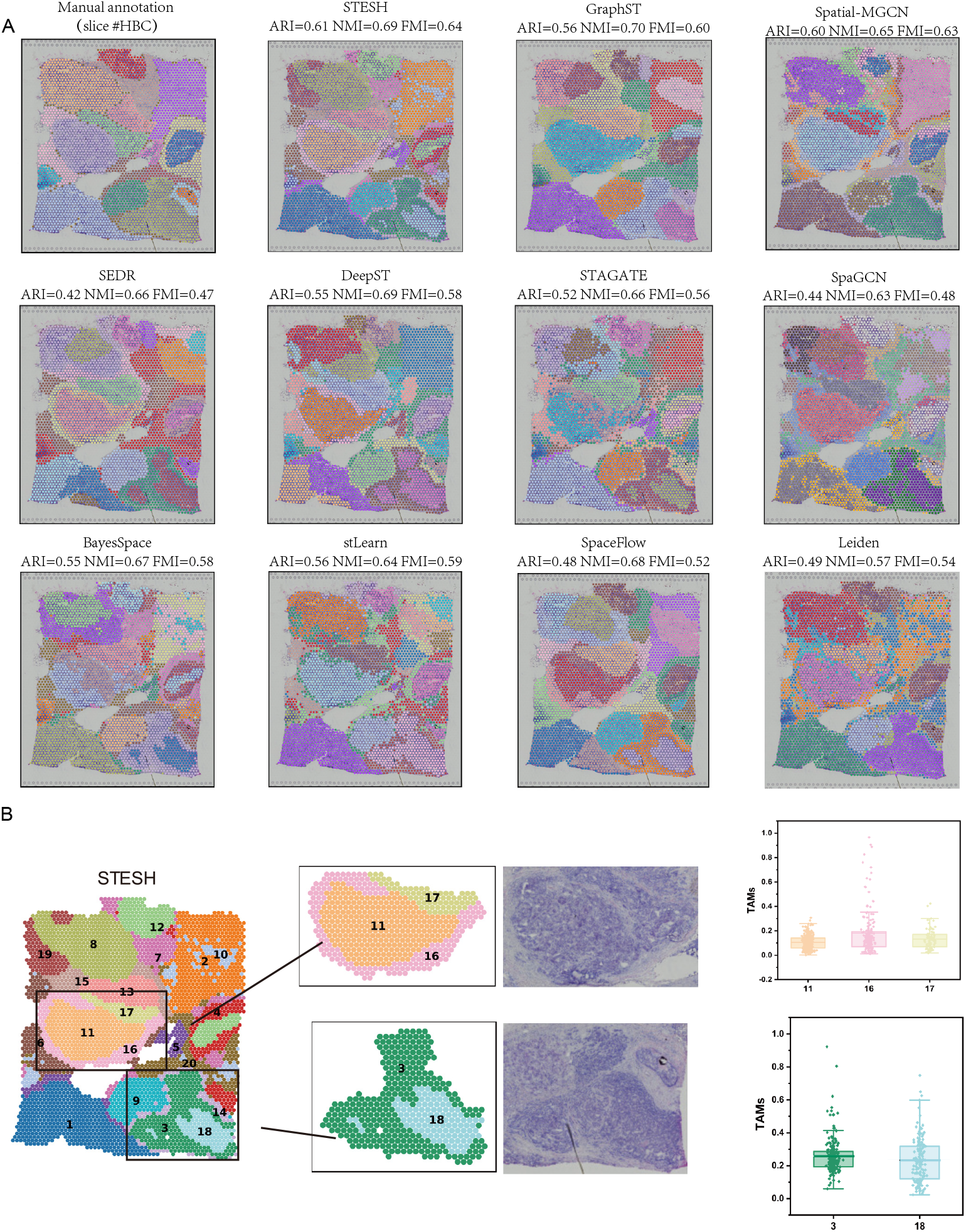
STESH improves spatial domain recognition in human breast cancer tissue. **A**, Identification of spatial domains by manual annotation, STESH, GraphST, Spatial-MGCN, SEDR, DeepST, STAGATE, SpaGCN, BayesSpace, stLearn, spaceFLOW, and Leiden. **B**, Visualization of cluster 11, 16, 17, 3 and 18 and their percentage of tumor-associated macrophages.

Cancer tissues exhibit significantly greater heterogeneity compared to normal cortical tissues. Accurately characterizing this complexity based solely on morphological features is challenging. However, spatial transcriptomics clustering methods that integrate gene expression data can uncover tumor heterogeneity missed by manual annotation. For instance, STESH, DeepST, and SEDR identified IDC_3 as comprising an outer “ring” and an inner core. Similarly, IDC_2 was subdivided by STESH, stLearn, DeepST, STAGATE, and SEDR, while DCIS/LCIS1 was split by STESH, STAGATE, and SEDR. To explore the biological significance of these classifications, we performed deconvolution analysis using Scanpy, aligning spatial data with scRNA-seq reference datasets^19^. We found that the percentage of tumor-associated macrophages (TAMs) of the outer ring is significantly higher than the inner core. TAM plays multi-function roles in cancer initiation and promotion, immune regulation, metastasis, and angiogenesis, such as providing essential support on tumor progression and metastasis^20^. Furthermore, the outer ring is also heterogeneous. STESH split the outer ring into clusters 16 and 17 (Figure 4b). Cluster 16 is directly adjacent to healthy tissue and had a higher TAM percentage, while cluster 17 is adjacent to another tumor tissue and had a similar TAM percentage to cluster 11.

### Benchmarking STESH on Stereo-seq dataset

New high-resolution spatial transcriptomics methods such as Stereo-Seq and Seq-Scope^21^ provided new opportunities for spatial transcriptomics tools. STESH on a Stereo-Seq mouse olfactory bulb dataset (19109 spots) is evaluated against two fine methods (STAGATE and SEDR) in previous benchmarking and one popular method (Leiden, Figure 4B). The histological image of the mouse olfactory bulb is briefly divided in seven layers, being the olfactory nerve layer (ONL), glomerular layer (GL), external plexiform layer (EPL), mitral cell layer (MCL), internal plexiform layer (IPL), granule cell layer (GCL), and rostral migratory stream (RMS, Figure 4A). Since there is no manual annotation for each spot, quantitative evaluation of the method performance by ARI, NMI or FMI is not achieved. Visually, all four methods reconstructed the biological structure of the mouse olfactory bulb layers (Figure 4C). STESH clustered this dataset into seven circular regions, which were similar to the marker genes for each layer (Figure 4D). In the UMAP and PAGA trajectory plots, SEDR, STAGATE, and Leiden exhibited network-like topological structures, indicating erroneous connections between clusters (Figure 5E). In contrast, STESH demonstrated an almost linear topological structure, precisely capturing the sequential development from the ONL to the RMS. These results highlight STESH’s superior clustering accuracy.

**Figure 5.**
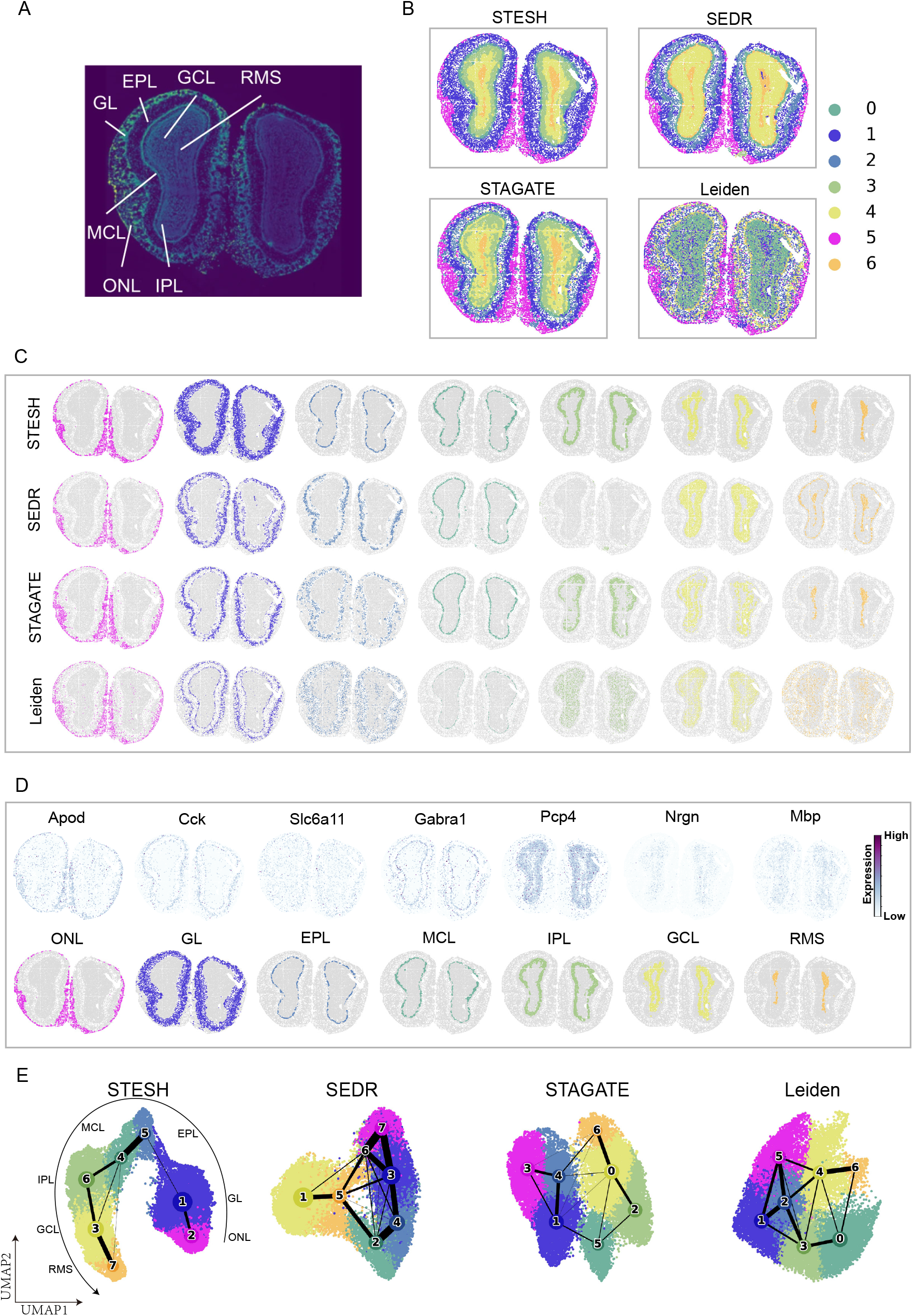
Application of STESH on Stereo-seq dataset. **A**, Laminar organization of a DAPI-stained mouse olfactory bulb. **B**, Clustering results from STESH, SEDR, STAGATE, and Leiden on the olfactory bulb Stereo-seq data. **C**, Visualization of individual identified clusters. **D**, Marker genes and predicted layers from STESH. **E**, Trajectory inference and UMAP visualization.

## Conclusion and Discussion

In this work, we introduce STESH, a novel method for processing spatial transcriptomics data to generate low-dimensional latent embeddings. STESH integrates spatial, RNA expression, and histological data using a multi-view graph convolutional network. Specifically, it constructs spatial adjacency matrices, feature adjacency matrices, and histological adjacency matrices, each based on different similarity metrics derived from tissue slice images, gene expression data, and spatial location information. By combining the gene expression matrix with the adjacency matrix, STESH builds spatial, feature, and histological graphs. The method employs a multi-view graph convolutional model to learn specific embeddings for each of these graphs. An attention mechanism is designed to adaptively learn the importance of each embedding, which is used to generate the final low-dimensional embedding. Clustering is performed based on learned embeddings to achieve spatial domain recognition, integrating multi-modal information in spatial transcriptomics. STESH enhances downstream analyses such as spatial clustering, UMAP visualization, and trajectory inference, offering new platforms and tools for spatial transcriptomics research. It addresses the limitations of existing methods in effectively utilizing spatial information, matching high-resolution histological images, and improving the accuracy of spatial domain recognition. It is a better way to precisely analyze spatial domains with similar gene expression and patterns in situ histology.

Accurate identification of spatial domains is fundamental for describing genomic heterogeneity and cell interactions, as well as for various downstream tasks in spatial transcriptomics analysis. Existing methods have limitations to effectively utilize spatial information and match high-resolution histological images. The strength of STESH lies in its deep integration of tissue image information. Instead of simply overlaying tissue image information upon gene expression data and spatial location information, morphological dependencies are constructed between spots using tissue images. This relationship is continuously and deeply considered at every stage of model training, achieving a greater level of data fusion.

## Method

### Morphological Feature Extraction

Image segmentation of tissue slice images is performed by determining the center of image blocks based on the actual coordinates of the spots in the slices. The tissue slice images are cropped into square image blocks of size 224×224 pixels. Morphological feature extraction from tissue slice images is carried out using a pre-trained ResNet50 model as a feature extractor, with the cropped image blocks as input, to extract the morphological information from the tissue slice images.

### Construction of Multi-Dimensional Adjacency Matrices

By calculating different similarity metrics, the spatial adjacency matrix A_*S*_, expression adjacency matrix A_f_, and morphological adjacency matrix A_m_ are constructed. These matrices, along with the gene expression matrix, are used for subsequent graph construction.

The spatial adjacency matrix is used to represent the spatial relationships between spots in the tissue. By calculating the physical distance between spots, it is possible to determine which cells are neighbors. This helps the subsequent model learn the gene expression patterns of spatially adjacent cells. The calculation is as follows:

First, the Euclidean distance *ds* between each spot and all other spots is computed based on the spatial location of the spots to measure spatial similarity. To define their adjacency relationships, a pre-defined radius r is used. The spatial adjacency matrix S between two spots is calculated by combining the Euclidean distance with the predefined radius r. The specific construction method is as follows: for a given spot, if the distance between the centers of two spots is smaller than the calculated radius r, they are considered adjacent. The corresponding position in the adjacency matrix is set to 1; otherwise, it is set to 0. The calculation formula is as follows:

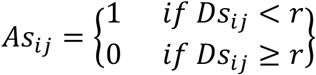

The expression adjacency matrix reflects the connectivity between spots at the gene expression level and is an important component of the expression graph. The calculation method is as follows: the cosine distance d_f_ is used to measure gene expression similarity, revealing the underlying structure of gene expression. For a given spot i and spot j, assuming their gene expressions are x_i_ and x_j_, the cosine distance is calculated using the formula:

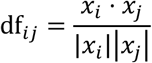

To better define gene expression similarity, a k-nearest neighbor graph is constructed for the gene expression matrix, referred to as the expression graph. This graph is represented by the expression adjacency matrix A_f_ for N spots, where the adjacency matrix is calculated based on the cosine distance d_f_. The top k most similar gene expression spots are defined as neighbors. The specific calculation method is as follows: for a given spot i, if spot j is a neighbor of spot i, then the corresponding position in the adjacency matrix is set to 1; otherwise, it is set to 0.

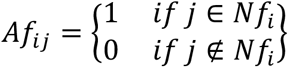

### Morphological Adjacency Matrix

The morphological adjacency matrix A_m_ reflects the relationship between cells or tissue regions based on their morphological features.

First, the tissue images are segmented based on the coordinates of each spot, and features are extracted from the images using a pre-trained convolutional neural network (CNN), which are then used as the morphological feature vector for each spot. Since the feature vectors extracted by the pre-trained convolutional neural network are high-dimensional, Principal Component Analysis (PCA) is used to select the top 50 features as the latent morphological feature representation M for each spot. Finally, for a given spot i and spot j, the Pearson correlation *dm* between the morphological latent features is calculated using the formula:

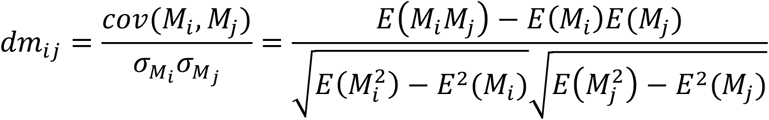

The top k most similar image blocks for each spot are identified as neighbors. for a given spot i, if spot j is a neighbor of spot i, then the corresponding position in the adjacency matrix is set to 1; otherwise, it is set to 0. The construction formula is the same as for the expression adjacency matrix.

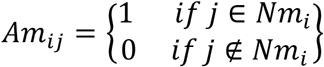

### Graph construction

Graph construction mainly includes the creation of the expression graph, spatial graph, and morphological graph.

The expression graph is based on the cosine distance d_f_ between each spot to measure gene expression similarity. These similarity metrics reflect the degree of similarity in gene expression between different spots, which is the key basis for constructing the expression graph. Based on the calculated gene expression similarity, the top k nearest neighbors for each spot are selected, and an adjacency matrix A_f_ representing gene expression similarity is constructed. Finally, the gene expression data is used as the node attribute feature matrix X and combined with the adjacency matrix, the complete expression graph G_f_(A_f_, X) is constructed.

The spatial graph is primarily based on spatial location information. By calculating the Euclidean distance between each spot in the tissue slice, spatial similarity is measured, and an adjacency matrix that reflects spatial similarity is constructed. Based on the computed Euclidean distance d_s_ and a pre-defined radius r, the adjacency matrix A_s_ is constructed. The gene expression feature matrix X is used as the node attribute feature matrix. The spatial graph G_s_(A_s_, X) is constructed using the spatial similarity adjacency matrix A_s_ and the node attribute feature matrix X.

The morphological graph is constructed based on the morphological information extracted from the tissue slices. The Pearson correlation d_m_ between the image blocks corresponding to each spot is calculated to measure morphological similarity. Using the computed Pearson correlation 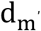 the top k nearest neighbors for each spot are selected, and an adjacency matrix A_m_ representing morphological similarity is constructed. The gene expression feature matrix X is used as the node feature matrix. The morphological graph G_m_(A_m_, X) is constructed using the morphological similarity adjacency matrix A_m_ and the node feature matrix X.

### Multi-View Graph Convolutional Autoencoder

The Multi-View Graph Convolutional Autoencoder (MCGCN) consists of spatial convolution modules, feature convolution modules, morphological convolution modules, and collaborative convolution modules. The method for generating low-dimensional embeddings is as follows:

1. The spatial convolution module performs convolution operations on the spatial graph and applies the following hierarchical propagation rule to generate low-dimensional embeddings E_s_:

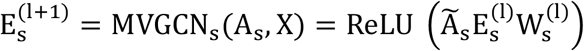

Where: 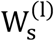 is the weight parameter of the l-th layer in the spatial convolution module; 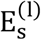 is the low-dimensional embedding generated by the l-th layer in the spatial convolution module; ReLU represents the ReLU activation function. Ã_s_is the symmetric normalized adjacency matrix in the spatial graph, and its calculation method is shown below:

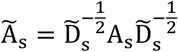

Where 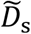 represents the degree matrix of A_s_
2. The feature convolution module performs convolution operations on the expression graph and applies the following hierarchical propagation rule to generate low-dimensional embeddings E_f_:

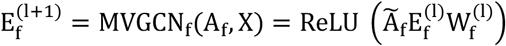

Where: 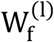 is the weight parameter of the l-th layer in the feature convolution module; 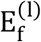 is the low-dimensional embedding generated by the l-th layer in the feature convolution module; Ã_f_ is the symmetric normalized adjacency matrix in the expression graph, and its calculation method is the same as for the spatial graph.
3. The morphological convolution module performs convolution operations on the morphological graph and applies the following hierarchical propagation rule to generate low-dimensional embeddings E_m_:

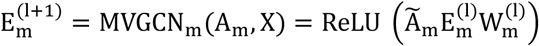

Where: 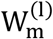 is the weight parameter of the l-th layer in the morphological convolution module; 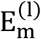 is the low-dimensional embedding generated by the l-th layer in the morphological convolution module; Ã_m_ is the symmetric normalized adjacency matrix in the morphological graph, and its calculation method is the same as for the spatial graph.
4. Since gene expression has a certain correlation with spatial distribution and histological information, a collaborative convolution module is introduced for collaborative convolution of three graphs to extract the collaborative embedding E_cm_, E_cs_, and E_cf_ for histological, spatial, and expression graph, respectively.

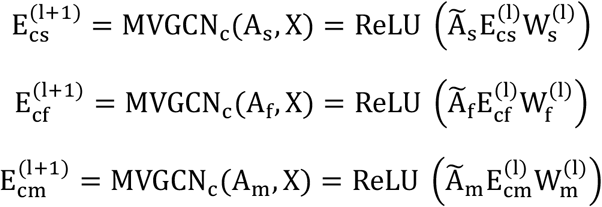

Through the above calculated 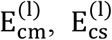 and 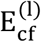, the collaborative embedding 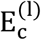 is defined as:

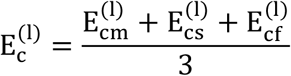

In order to learn a more consistent representation, the consistency is measured bycomparing the difference in the covariance matrices between E_cm_, E_cs_, and E_cf_. Define the consistency constraint loss L_con_ as:

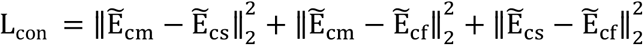

Where: - 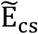 is the embedding extracted from the spatial graph, 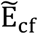 is the embedding extracted from the expression graph, 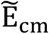 is the embedding extracted from the morphological graph.

### Attention mechanism

An attention mechanism is introduced to adaptively learn *E*_*m*_, *E*_*s*_, *E*_*f*_, and *E*_*c*_ generated by the multi-view graph convolution autoencoder, to generate the corresponding weight parameters *ω*_*m*_, *ω*_*s*_, *ω*_*f*_, and *ω*_*c*_, and to generate the final low-dimensional embedding *E*_*final*_ through the weight parameters.

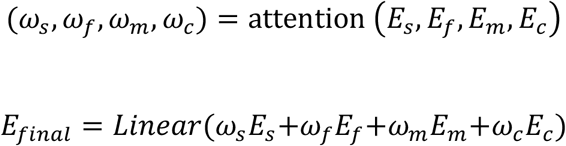

### Negative binomial decoder

The negative binomial decoder used the negative binomial distribution to model the characteristics of the data. When reconstructing the gene expression matrix, it accounts for the discrete and variable nature of gene expression data, enabling it to capture the complex global patterns present in ST data. Its composition is as follows.

Firstly, an intermediate layer containing a linear layer and a batch normalization layer is defined, which is used to map the low-dimensional latent embedding *E*_*final*_ (the encoder output) to a higher-dimensional space for extracting higher-level features. The ReLU activation function is used to introduce nonlinearity. Secondly, two linear layers are introduced to map the output of the middle layer to the original dimensions, respectively. Dispersion θ and mean µ of the distribution are obtained. Both the dispersion. and the mean are processed by different activation functions to ensure that their values are within range. This design helps the model to better fit the real data. Specifically, for a given gene expression matrix *X*, assuming that it conforms to the negative binomial distribution, the probability distribution of gene expression is defined as follows:

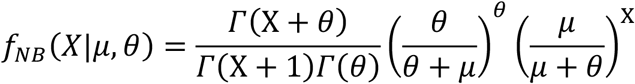

Where *μ* and *θ* are calculated by the decoder, representing the mean and dispersion, respectively. *Γ* represents the gamma function. In order to minimize the difference between the predicted value and the true value, the negative log-likelihood estimation is used to reconstruct the loss of the original gene *L*_NB_rec_:

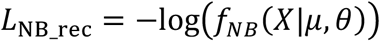

### Loss function

To better capture the structural information of the graph, regularization constraints are based on the spatial adjacency matrix, spatial negative sampling matrix, histological adjacency matrix, and histological negative sampling matrix. Taking into account the influence of multiple graph structures, the model can learn the structural information of the graph more comprehensively by comparing the cosine similarity between nodes. The regularization constraint loss 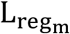 of the histological graph is defined as follows:

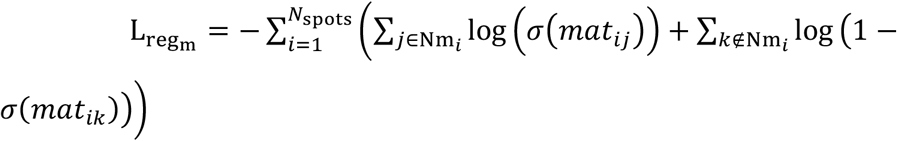

Where: Nm_*i*_ is the histological neighbor set of the spot *i* and *mat*_*ij*_ is based on the cosine similarity matrix of the learned latent representation *E*_*final*_. The loss function contains two parts the histological adjacency matrix positive sample loss is to encourage the embedding vectors of histological adjacent nodes to be closer in the embedding space to learn the local structure in the histological graph, histological adjacency matrix negative sample loss is to encourage the embedding vectors of histological non-adjacent nodes to be further apart in the embedding space to avoid noise and overfitting in the learned graph.

Similarly, the regularization constraint loss 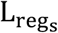 of the spatial graph is defined as follows:

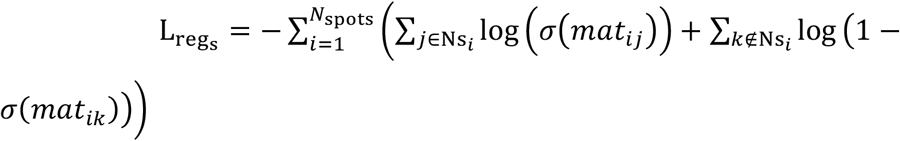

Taking into account the regularization constraint loss 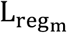 of the histological graph and the regularization constraint loss 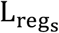 of the spatial graph, the overall regularization constraint loss 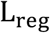 is defined as follows:

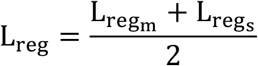

In summary, the STESH model consists of the multi-view graph convolution encoder and the negative binomial decoder. The total loss function *L* is composed of the reconstruction loss *L*_*NB*_*rec*_ of the original gene, the consistency constraint loss *L*_*con*_, and the regularization constraint loss *L*_*re*g_.

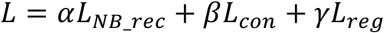

### Downstream analysis

The learned latent representation can be applied to downstream analysis tasks. To cluster the spatial data, STESH uses mclust() in R. To visualize it in two dimension, STESH uses sc.pl.umap() function in Scanpy. To infer the trajectory, STESH uses sc.tl.paga() function in Scanpy.

### Data Preprocessing

For all datasets, to reduce technical noise in ST data, we first remove spots outside the main tissue regions. Then, we use the SCANPY package to filter out genes with low expression or low variance from the raw gene expression data, selecting the top 3000 highly variable genes. Finally, we normalize the data using a scaling factor. The normalization function is defined as follows:

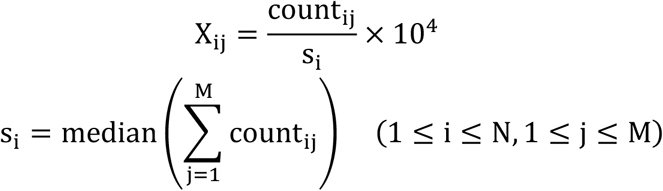

where X_ij_ denotes the original expression value of the j-th gene in the i-th spot.

### Parameter Settings

The entire model is implemented using PyTorch 2.6.0. In the experiments of this model, two graph convolution layers are used for the three views, where the output dimension of the first convolution layer is set to 128, and the output dimension of the second convolution layer is set to 64. To optimize the model, the Adam optimizer is employed with a learning rate of lr = 0.001. The weight parameter values are set to α=1,β=10and γ=0.1. For all baseline methods, we use the default parameters from the original paper.All experiments are conducted on a server with an NVIDIA GeForce RTX 3090 GPU running Ubuntu 20.04.5.

## Contributions

HZ: software, visualization, writing—original draft. JC: methodology, software, formal analysis. YS: visualization. PH: visualization. JingLiu: visualization. JiujuLuo: visualization. HY: software. YR: writing—review and editing. XZ: supervision. ZC: supervision. KWW: supervision, writing—review and editing. HS: conceptualization, funding acquisition.

## Code Availability

The source code has been publicly released under the MIT open-source license, which complies with the requirements of the Open Source Initiative. The source code of STESH is deposited as a Python package on Zenodo (https://doi.org/10.5281/zenodo.14954828), and it is also available on GitHub (https://github.com/haojingshao/STESH).

## Data Availability

The datasets used in this paper can be downloaded from the following websites. (1) The LIBD human dorsolateral prefrontal cortex (DLPFC) dataset (http://spatial.libd.org/spatialLIBD); (2) 10x Visium spatial transcriptomics dataset of human breast cancer(https://support.10xgenomics.com/spatial-gene-expression/datasets/1.1.0/V1_Breast_Cancer_Block_A_Section_1);and(3)the processed Stereo-seq dataset from mouse olfactory bulb tissue (https://github.com/JinmiaoChenLab/).

